# Murine modeling of menstruation identifies immune correlates of protection during *Chlamydia muridarum* challenge

**DOI:** 10.1101/2024.05.21.595090

**Authors:** Laurel A. Lawrence, Paola Vidal, Richa S. Varughese, Zheng-Rong Tiger Li, Thien Duy Chen, Steven C. Tuske, Ariana R. Jimenez, Anice C. Lowen, William M. Shafer, Alison Swaims-Kohlmeier

## Abstract

The menstrual cycle influences the risk of acquiring sexually transmitted infections (STIs), including *Chlamydia trachomatis* (*C. trachomatis*), although the underlying immune contributions are poorly defined. A mouse model simulating the immune-mediated process of menstruation could provide valuable insights into tissue-specific determinants of protection against chlamydial infection within the cervicovaginal and uterine mucosae comprising the female reproductive tract (FRT). Here, we used the pseudopregnancy approach in naïve C57Bl/6 mice and performed vaginal challenge with *Chlamydia muridarum* (*C. muridarum*) at decidualization, endometrial tissue remodeling, or uterine repair. This strategy identified that the time frame comprising uterine repair correlated with robust infection and greater bacterial burden as compared with mice on hormonal contraception, while challenges during endometrial remodeling were least likely to result in a productive infection. By comparing the infection site at early time points following chlamydial challenge, we found that a greater abundance of innate effector populations and proinflammatory signaling, including IFNγ correlated with protection. FRT immune profiling in uninfected mice over pseudopregnancy or in pig-tailed macaques over the menstrual cycle identified NK cell infiltration into the cervicovaginal tissues and lumen over the course of endometrial remodeling. Notably, NK cell depletion over this time frame reversed protection, with mice now productively infected with *C. muridarum* following challenge. This study shows that the pseudopregnancy murine menstruation model recapitulates immune changes in the FRT as a result of endometrial remodeling and identifies NK cell localization at the FRT as essential for immune protection against primary *C. muridarum* infection.

**Author Summary:** Although the vast majority of women and adolescent girls of reproductive age experience menstruation, we have little insight into how this tissue remodeling process alters mucosal immune defenses against infection by genitourinary pathogens. In this study, we used a murine model of menstruation to investigate how endometrial shedding and repair alters the immune landscape in the female reproductive tract (FRT) to influence chlamydial infections. Using this approach, we identified that endometrial remodeling regulates a substantial pro-inflammatory immune response, including NK cell recruitment into the cervicovaginal tissues, and we further confirmed this phenomenon is occurring in a naturally menstruating species. The localization of NK cells in the FRT at the time of challenge was determined to be responsible for rapid immune protection that reduced *C. muridarum* burden, as experimental depletion of these cells over this timeframe now led to productive infections. Taken together, this study identifies that murine models of menstruation can be a valuable tool for investigating how the menstrual cycle modulates immune homeostasis and for identifying ways to strengthen mucosal immune defenses against genitourinary pathogens in women.

## Introduction

*Chlamydia Trachomatis* (*C. trachomatis*) infections spread through sexual contact and can result in severe diseases in women and congenitally infected newborns. *C. trachomatis* is the causative agent of one of the most common and costliest bacterial sexually transmitted infections (STIs) globally, with the majority of infections occurring in women and adolescent girls of reproductive age [1]. Although chlamydial infections remain an urgent global health issue, there is currently no vaccine that can protect against *C. trachomatis*. Notably, reinfections are common, which increases the likelihood of developing severe diseases, including pelvic inflammatory disease (PID), stillbirths, infertility, and an increased risk of acquiring more severe secondary infections such as those caused by *Neisseria gonorrhoeae* (*N. gonorrhoeae*) and human immunodeficiency virus type-1 (HIV) [2]. Although immune cells positioned within mucosal barrier sites, such as the cervicovaginal and uterine mucosae of the female reproductive tract (FRT), can provide an immediate effector response against invading pathogens due to their proximity at an infection site [3–5], these contributions to *C. trachomatis* infections of the FRT are unclear. Thus, greater insights into how the FRT tissue environments regulate cellular immune barrier defense against chlamydial infections are essential for informing prevention efforts.

Globally, the vast majority of women (in addition to some transgender men and gender non-conforming persons assigned female at birth) of reproductive age (15-49 years) experience periodic menstruation [6], and though dependent upon the immune system, the processes by which menstruation can influence protection against invading pathogens are poorly recognized. In regards to infection risk by the most prevalent bacterial STI pathogens in the U.S., including *C. trachomatis* and *N. gonorrhoeae,* it has been previously shown using animal modeling and human tissue samples that levels of the sex hormones progesterone and estrogen are associated with infection risk and the potency of immune effector responses against chlamydial infections [7–14]. However, we have limited insights into what role the menstrual cycle plays in determining these differences. A major challenge in studying how menstruation impacts immune defenses at barrier sites is a lack of model systems that menstruate [15], especially common laboratory animal models that are supported by immunologic and genetic approaches that could facilitate mechanistic investigations into the dynamics of FRT tissue-localized immune cell populations [16–22].

Previously, a minimally invasive strategy for inducing menstruation in the BALB/c strain of inbred laboratory mice was reported using the pseudopregnancy method [23]. In this approach, BALB/c female mice in estrus were mated with vasectomized males to induce uterine decidualization. At the time frame of implantation, sesame seed oil is injected into the endometrial environment, leading to a state of terminal differentiation by decidual cells. The subsequent decline in progesterone causes rapid deterioration of the endometrium, which prompts uterine remodeling and discharge in the mice, similar to the process of menstruation occurring in species that naturally undergo spontaneous decidualization, such as humans and pig-tailed macaques [24, 25].

To test whether the pseudopregnancy approach for inducing menstruation in mice might provide insights into tissue-specific immune determinates of *C. trachomatis* infection, we applied this method to the C57Bl/6 strain of inbred mice paired with vaginal challenge by *Chlamydia muridarum* (*C. muridarum)*, a murine strain of chlamydia which models lower FRT infection by *C. trachomatis* [26, 27]. This strategy showed that following the induction of pseudopregnancy, C57Bl/6 mice exhibited progesterone fluctuations in circulation with corresponding innate immune cell recruitment into both the uterine horns and cervicovaginal tissues followed by decidual discharge (*i.e.,* menses). By performing vaginal challenges with *C. muridarum* based on the time point of pseudopregnancy, we found that while challenges administered under conditions of uterine repair resulted in robust infections, whereas challenges administered during decidualization and endometrial remodeling were unlikely to result in a production infection. Immune profiling of the FRT tissues showed that endometrial remodeling was associated with increased IFNγ signaling and NK cell recruitment in the cervicovaginal tissues. To test whether this change occurs in naturally menstruating species, we used longitudinal measurements from female pig-tailed macaques of reproductive age and confirmed both increased IFNγ-associated signaling and NK cell infiltration at the cervicovaginal lumen during conditions of endometrial remodeling. Finally, to confirm the role of NK cells in early protection against primary *C. muridarum,* we depleted NK cells in mice during the time span of endometrial remodeling prior to vaginal challenge, which then resulted in productive chlamydial infections.

Taken together these data show that the menstrual cycle determines NK cell localization within the cervicovaginal mucosa, which plays an essential role in early immune protection against primary *C. muridarum* infection. Importantly, we demonstrate that the murine pseudopregnancy method for inducing menstruation is a valuable tool for investigating mucosal immune correlates of protection and risk against chlamydial infection and potentially for developing strategies that can strengthen mucosal immunity against genitourinary pathogens.

## Results

To investigate immune changes in the FRT occurring as a result of menstruation, we began by optimizing the murine pseudopregnancy approach to induce overt menstruation in C57Bl/6 mice. Female mice aged 6-12 weeks were mated with vasectomized males, and successful ejaculation was confirmed by the detection of a vaginal plug the following morning **(Figure 1A)**. Monitoring sex hormones in circulation over pseudopregnancy **(Figure 1B)**, we observed that progesterone levels sharply increased by day 4, and by day 6, progesterone levels in circulation had reached a peak. Progesterone withdrawal due to the absence of fertilization was detected on day 8, and by day 10, progesterone levels decreased to a range consistent with those detected on day 2 and in mice treated with the hormonal contraceptive medroxyprogesterone acetate (MPA) to control for reproductive cycling. Over the course of pseudopregnancy, the lowest levels of progesterone were detected on day 12. In contrast to progesterone, levels of estrogen generally remained in the range of those detected in MPA-treated mice and did not significantly change until day 14, at which point estrogen sharply increased **(Figure 1C)**. Next, to identify how pseudopregnancy impacted cellularity of the vaginal environment, we monitored changes by vaginal cytology **(Figure 1D)**. Microscopy of vaginal smears showed an increase in neutrophils during the time frame of endometrial remodeling (days 6-8), followed by the detection of red blood cells between days 10-11. On days 12-14, when estrogen levels peaked, we identified a predominant population of anucleated vaginal epithelial cells consistent with ovulation [28]. Using ALPHA-dri bedding, vaginal swabs, or visual inspection **(Figure 1E)**, we confirmed that mice were menstruating between days 10-11. Next, to measure uterine vascularization over pseudopregnancy, we performed intravital labeling of circulating leukocytes **(Figure 1F, G)** [16] prior to necropsy and compared the frequency of circulating cells in the uterine horns of pseudopregnant mice to mice treated with MPA or sesame seed oil control mice (mice receiving an intrauterine injection with sesame seed oil but not mated with vasectomized male mice) **(Figure 1H)**. This approach showed that starting on day 4, the mean levels of vascular cells began to increase, reaching the highest frequency, on average about 40% of the overall leukocyte population, on day 8. By day 12, the frequency of circulating cells was once again decreased into the range detected on day 2 of pseudopregnancy, similar to MPA and sesame seed oil control mice. Overall, these data showed that the pseudopregnancy approach was reciprocated in the C57Bl/6 mice and identified key time points of endocrine-regulated endometrial remodeling and menstruation.

**Figure 1.**
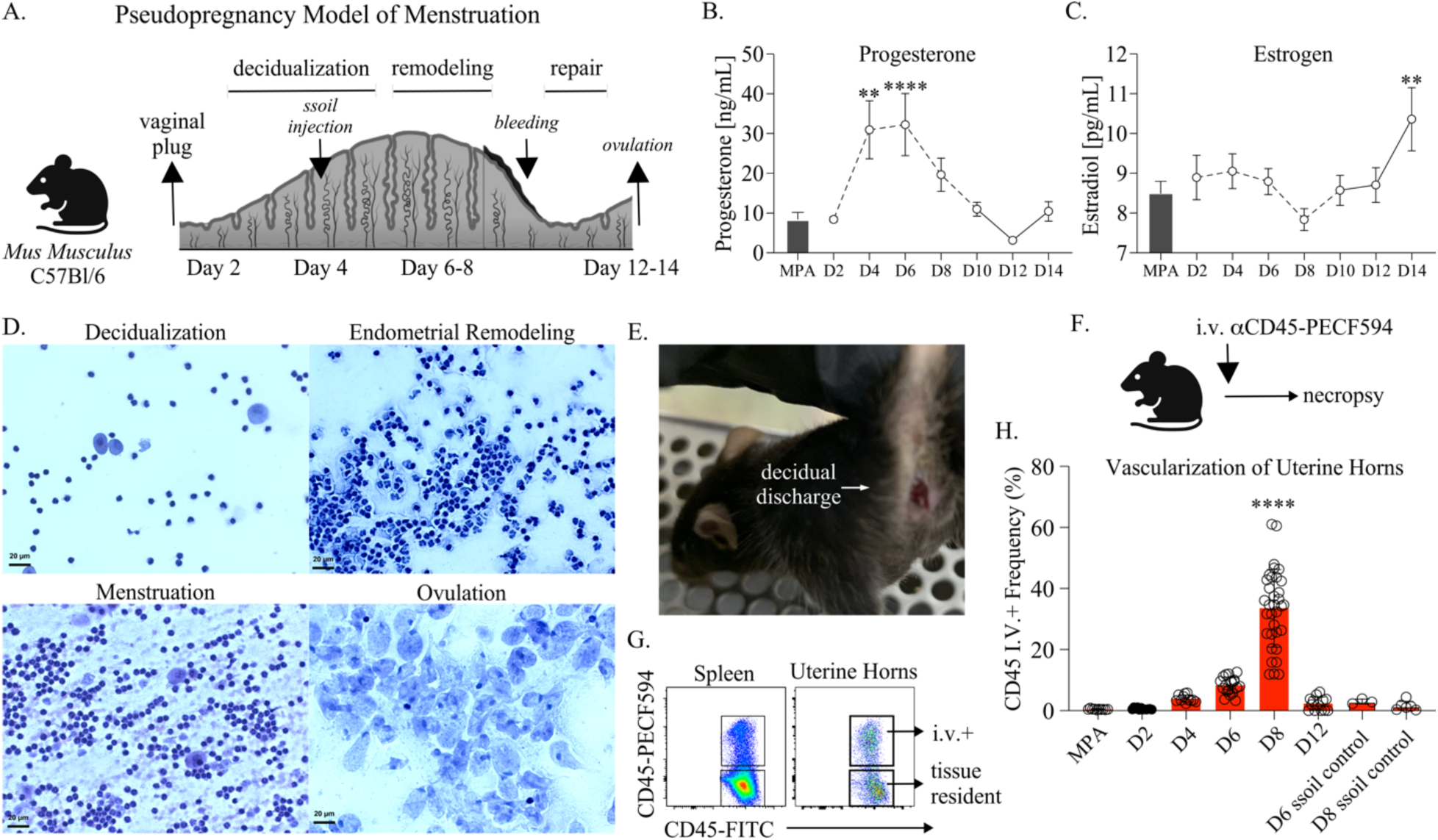
**(A)**. Depiction of the pseudopregnancy approach for inducing menstruation in C57Bl/6 mice with the time frame of major endometrial changes emphasized. **(B)**. A mean symbol graph with the standard error of means (SEM) depicting Progesterone or **(C).** Estrogen levels measured from blood plasma over indicated time points of pseudopregnancy and plotted as concentration. Progesterone and Estrogen levels from mice administered Medroxyprogesterone acetate (MPA) are shown as a bar graph with SEM for each comparison. **(D)**. Vaginal cytology over pseudopregnancy at indicated time points. Vaginal smears are stained using Hematoxylin and Eosin (H&E) and then visualized using microscopy at 40x magnification. **(E)**. A photo of a menstruating C57Bl/6 mice following the induction of pseudopregnancy (day 10). **(F)**. A schematic depicting the intravenous (IV) labeling approach for distinguishing leukocytes in circulation performed prior to euthanasia and necropsy. **(G)**. Cell flow plots from the spleen and uterine horns of a representative animal at day 8 of pseudopregnancy, illustrating the approach for distinguishing tissue-resident or circulating leukocyte measurements following IV labeling. Viable, singlet leukocytes are distinguished by the expression of IV-labeled CD45. **(H)**. The frequency of IV+ leukocytes from uterine horns over the indicated time points of pseudopregnancy and compared with sesame seed oil (ssoil) injection control mice or mice treated with MPA. Each open circle represents an individual mouse **(B, C)**. Models used to evaluate a mean deviation were fit using one-sample t-tests. A minimum of 6 mice were measured at each time point. **(H)**. Models used to compare a difference of means were fit using multiple comparisons: **(B, C, H)**. p-values with q-values ≤ 0.05 are shown. *p≤0.05, **p<0.01, ***p<0.001, ****p<0.0001. **(A, F)**. Created using BioRender.com.

Next, to elucidate changes in the immune landscape over endometrial remodeling and menstruation in the FRT, we performed longitudinal cellular profiling of innate immune cell populations previously identified as important for protection against primary chlamydial infections; neutrophils, macrophage, and NK cells **(Figure 2)** [29–31]. To identify potential differences in FRT tissue compartments, the FRT tissues were first distinguished from luminal cells collected by cervicovaginal lavage (CVL), followed by dissection of cervicovaginal tissues (or lower FRT, LFRT) from the uterine horns. Single-cell suspensions were then assessed for immune cell populations using flow cytometry **(Figure 2A)**. By quantifying tissue-resident leukocyte yields over pseudopregnancy **(Figure 2B and Supplemental Figure 1)**, we identified that neutrophil populations increased within the uterine tissues at the time point progesterone levels in blood peaked (day 6 of pseudopregnancy) and were entering into a state of withdrawal, similar to previous reports [32, 33]. While endometrial neutrophils have been previously implicated in facilitating endometrial shedding and repair, we also observed neutrophil infiltration into the LFRT. However, at the luminal surface (CVL), neutrophil numbers were not changed at day 6 compared with day 4, although these day 6 levels were higher compared to the time frame of uterine repair and ovulation (day 12) and from mice that were treated with MPA. Corresponding to neutrophil changes, we identified a similar trend of increased macrophage populations at day 6 throughout the FRT, which have also been identified as important contributors to endometrial tissue breakdown and repair during menstruation [34]. Next we examined NK cell populations, which in the uterus are thought to play a critical role in implantation and have been shown to increase during the luteal phase when blood progesterone levels peak [35, 36]. We observed similar NK cell increases throughout the FRT on day 6 as compared with day 4; however, while neutrophils, macrophage, and NK cell numbers began to contract in the uterine horns on day 8, the luminal and LFRT NK cell numbers were sustained before ultimately decreasing by day 12. By comparing NK cells over pseudopregnancy with MPA treatment, we found that while the number of NK cells in the LFRT tissues at days 6-8 were within a similar range, luminal NK cell numbers were significantly greater on days 6-8 of pseudopregnancy as compared with MPA treatment.

**Figure 2.**
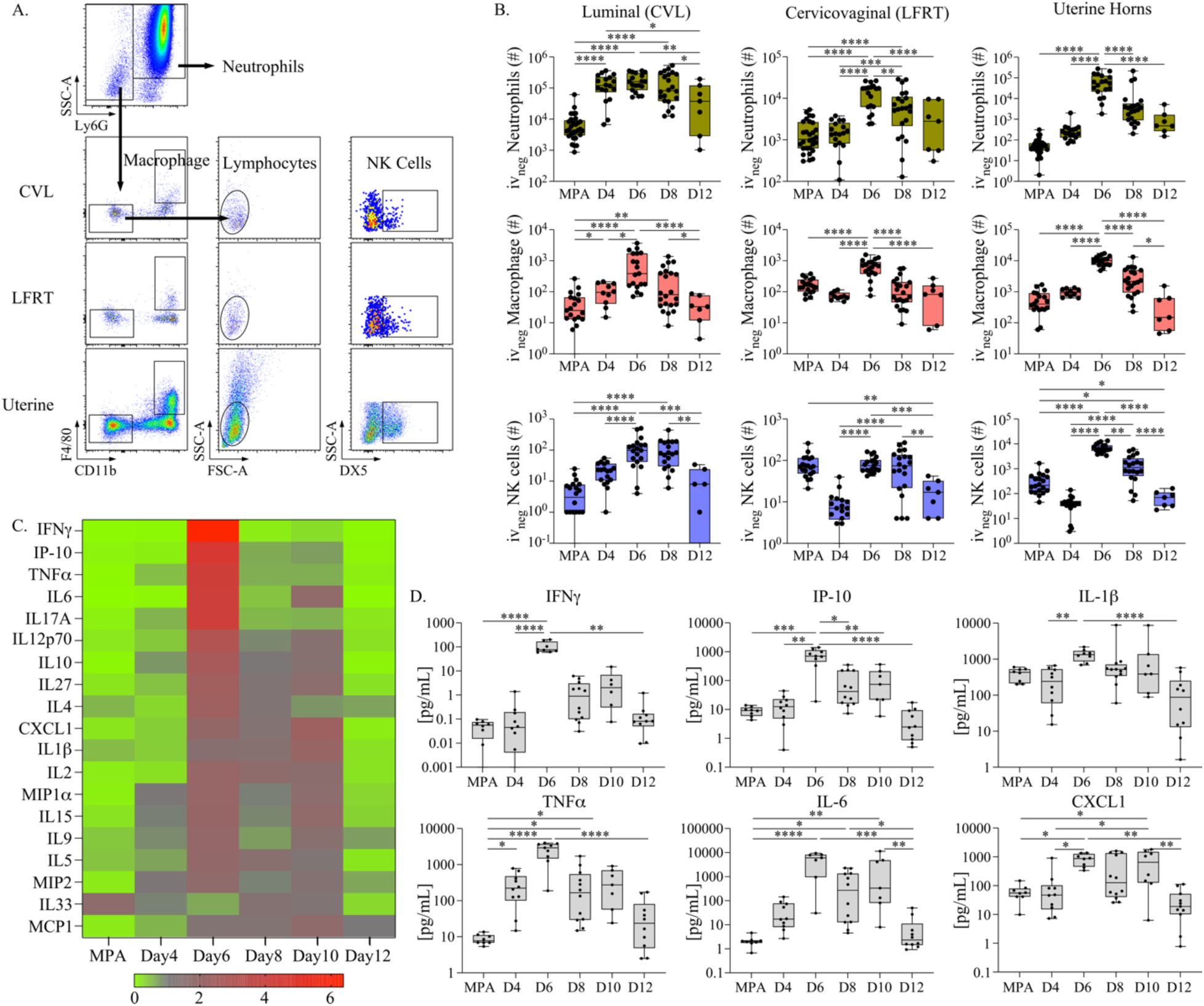
**(A)**. Flow cytometry cell gating strategy for measuring innate immune cells from anatomic compartments of the FRT. Viable singlet tissue-resident leukocytes are discriminated by the expression of Ly6G and side scatter characteristics from CVL, cervicovaginal tissue, and the uterine horns. The remaining populations are then measured for macrophage based on CD11b and F4/80 expression, and then NK cells are measured from CD11b and F4/80 negative lymphocytes (based on size and granularity characteristics) followed by DX5 expression. **(B).** The total yield of indicated immune cell populations from FRT tissue sites is plotted as bar and whiskers graphs over pseudopregnancy and compared with mice administered MPA as a control. **(C).** A heat map depicting the fold change in cytokines and chemokines measured from CVL over pseudopregnancy or MPA and ordered according to the greatest fold increase (top to bottom). **(D).** The concentrations of indicated cytokines are plotted as bar and whiskers graphs over pseudopregnancy and compared with mice administered MPA as a control. **(B, D).** Models used to compare a difference of means were fit using multiple comparisons: p-values with q-values ≤0.05 are shown *p≤0.05, **p<0.01, ***p<0.001, ****p<0.0001.

We also evaluated soluble immune mediators in cervicovaginal secretions over pseudopregnancy by measuring 19 pro-inflammatory cytokines and chemokines from CVL supernatants **(Figure 2C and summarized in Supplemental Table 1)**. This showed that, corresponding to the increases in the cellular immune populations, the levels of most cytokines and chemokines exhibited sharp elevations at day 6 of pseudopregnancy, with the greatest increases in IFNγ and the IFNγ-induced protein, IP-10. Because leukocyte infiltration during endometrial remodeling has been previously linked with increased uterine TNFα, IL1β IL-8 (murine homolog: CXCL1) and IL-6 production [32, 37], we specifically compared those CVL concentrations detected over pseudopregnancy in addition to IFNγ signaling **(Figure 2D)**. This analysis also showed sharp increases in the concentrations of these cytokines and chemokines at day 6 of pseudopregnancy, which by day 12 had decreased to levels detected within the ranges of day 4 and MPA. Although less well characterized in the context of the menstrual cycle, to our knowledge, IFNγ signaling has been previously identified as important for facilitating implantation and pregnancy [38–40] and is predominantly produced by NK cells, suggesting similar regulation by the menstrual cycle [41]. Thus, taken together, these data show that the LFRT and lumen also experience proinflammatory changes similar to the endometrium over pseudopregnancy. The only exception to these observations was the discovery of a sustained elevation of LFRT and luminal NK cells during the time frame of progesterone withdrawal.

As the changes observed in NK cells and IFNγ signaling in the cervicovaginal environment over pseudopregnancy were, to our knowledge, less defined in menstruating species, we tested whether these discoveries were translationally relevant by measuring these properties in pig-tailed macaques **(Figure 3A)**. To perform a longitudinal analysis over the menstrual cycle, 6 female pig-tailed macaques of reproductive age were sampled bi-weekly for blood to measure estrogen and progesterone and weekly for CVL collection for a period of 9 weeks. To stratify immune measurements, estrogen peaks (representing ovulation) and observed menstruation were used to determine cycle phases relative to the average cycle length of a pig-tailed macaque **(Figure 3B)**. Using leukocyte-enriched CVL cells compared with PBMC (peripheral blood mononuclear cells), NK cells were assessed by flow cytometry **(Figure 3C)**. To control for animal-to-animal variations, NK cell yields from each animal were calculated as a fold change. Luminal NK cell numbers at the luteal phase (peak progesterone) and late luteal phase (detection of progesterone withdrawal) in macaques were increased, consistent with sustained elevations observed during pseudopregnancy in the mice. Following these peaks in NK cell yields, levels then decreased prior to the onset of menstruation. Next, we evaluated IFNγ signaling by measuring the IFNγ-induced protein, IP-10, which is generally found at higher concentrations and, thus, was more likely to fall within the range of assay detection for NHP **(Figure 3D)**. The levels of CVL IP-10 also peaked at the luteal phase, matching peak progesterone, followed by a rapid decrease, similar to the kinetic trends observed in the mice. Taken together, these data show that the pig-tailed macaques exhibit sinusoidal patterns of NK cell recruitment and IFNγ signaling within the cervicovaginal tissue environment over the menstrual cycle, paralleling the murine pseudopregnancy approach for inducing menstruation.

**Figure 3.**
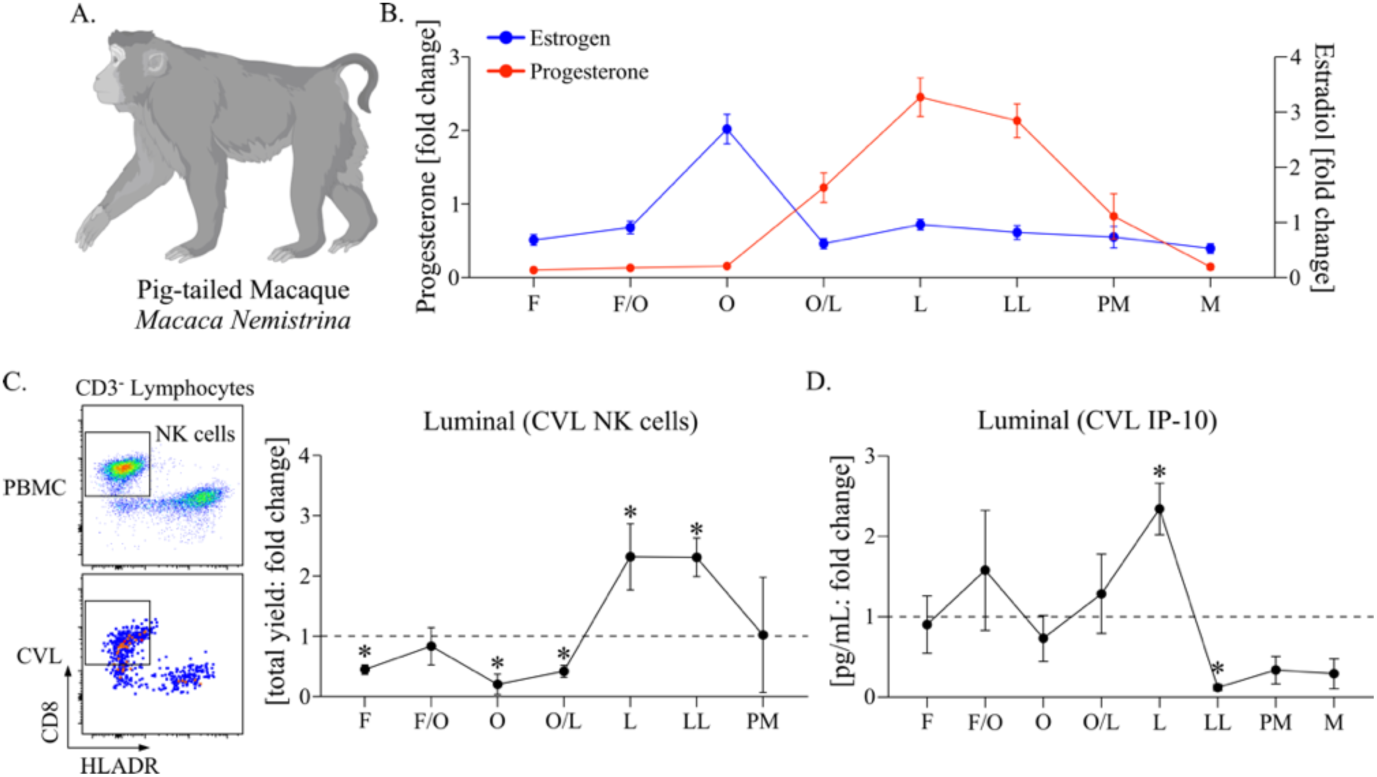
**(A)**. Cartoon of a pig-tailed macaque (*Macaca Nemistrina*). Figure created using BioRender.com. **(B).** A symbol line graph with SEM depicting the fold change in plasma levels of Progesterone (red symbols and lines) and Estrogen (Estradiol, blue symbols and lines) was measured longitudinally from 6 animals and stratified by cycle phase. **(C).** (Left panel) Flow dot plots depicting the strategy for measuring NK cells from PBMC (top) and CVL (bottom). Live, singlet CD45 expressing lymphocytes are first discriminated from granulocytes, myeloid cells, T cells, and B cells and then measured for HLADR negative and CD8 positive populations. (Right panel) a symbol line graph with SEM depicting the fold change in CVL NK cells measured longitudinally and stratified by cycle phase. Cells collected during time points of menstruation were considered contaminated by cells in blood circulation and are not shown. **(D).** A symbol line graph with SEM depicting the fold change in IP-10 measured from CVL supernatant **(B-D).** The cycle phases are identified as follows: Follicular phase (F), Follicular/Ovulation transition (F/O), Ovulation (O), Ovulation/Luteal transition (O/L), Luteal phase (L), Late Luteal phase (LL), Pre-menstruation (PM), and Menstruation (M). **(C, D).** Models used to evaluate fold change (against a value of 1) were fit using Wilcoxon rank sum tests. Median differences with p-values ≤ 0.05 are indicated by an asterisk.

To investigate how FRT immune dynamics over menstruation might influence the risk of chlamydial infection, we vaginally challenged mice with 1x10^5^ inclusion forming units (IFU) of *C. muridarum* at specific time points over pseudopregnancy **(Figure 4A)**. First, we measured chlamydia replication over the course of infection by comparing the time points spanning decidualization and endometrial remodeling at challenge (day 4, day 6, and day 8) with control mice administered MPA [42] **(Figure 4B)**. These data showed that, compared with MPA control mice, mice challenged at day 4, day 6, or day 8 of pseudopregnancy all exhibited little to no bacterial replication. Notably, at the peak of infection (7 days post-challenge), the levels of *C. muridarum* DNA were all significantly lower. The most pronounced difference was observed from challenges on day 8 of pseudopregnancy, which also presented significantly reduced replication at 3 days post-challenge. In contrast, *C. muridarum* challenge during a time point spanning menstruation (administered prior to the detection of menses), which is also when endometrial repair initiates (day 10), showed significantly increased levels of bacterial replication. The peak of infection showed a mean 1-log increase compared to MPA control mice and an overall mean increase in bacterial burden throughout the infection course **(Figure 4C)**. Overall, this approach showed that the timing of challenge over pseudopregnancy resulted in dramatic differences in the outcome of infection, typified by little to no bacterial replication detected from challenges prior to or during endometrial remodeling, but high bacterial replication detected when challenges were administered under conditions of menstruation/endometrial repair.

**Figure 4.**
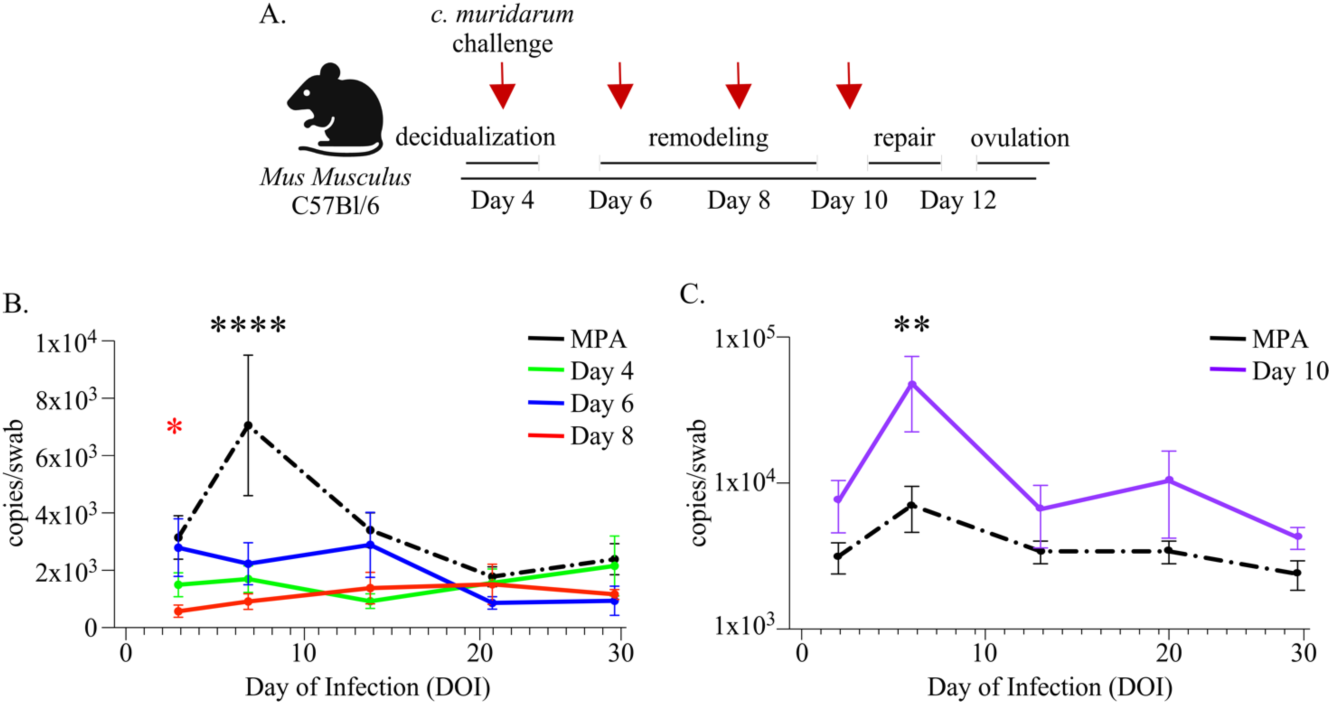
**(A)**. Schematic depiction of the vaginal *C. muridarum*challenge approach at time points of pseudopregnancy (indicated by red arrows). Created using BioRender.com. **(B, C).** Line graphs depicting the mean bacterial burden with SEM over the course of infection determined by ddPCR and compared with mice administered MPA prior to challenge (n=10, black dotted lines). The graphed data points based on the day of pseudopregnancy at challenge are comprised of 2 separate experiments for each group **(B).** The time of challenge over pseudopregnancy is indicated as day 4 challenge (n=8, green line), day 6 challenge (n=12, blue line), and day 8 challenge (n=10, red line). **(C).** The time of challenge over pseudopregnancy is indicated as day 10 (n=14, purple line). (B, C). Models used to compare a difference of means were fit using multiple comparisons: p-values with q-values ≤0.05 are shown *p≤0.05, **p<0.01, ***p<0.001, ****p<0.0001. (B). The red asterisk indicates a significant difference detected at day 3 post-challenge when comparing day 8 of pseudopregnancy at challenge with MPA.

To better understand the immune contributions to the differences observed with *C. muridarum* infection following vaginal challenge, we performed soluble and cellular immune profiling from the cervicovaginal tissues early in the infection course (3 days post-challenge) and compared these measurements by the timing of challenge over pseudopregnancy **(Figure 5)**. First, we measured cytokines and chemokines from cervicovaginal secretions and compared these levels between challenges occurring on day 8 of pseudopregnancy, when we detected the greatest decrease in overall bacterial burden, with challenges on day 10 of pseudopregnancy, when we detected the greatest increase in overall bacterial burdens **(Figure 5A)**. This analysis showed that, in general, animals challenged with chlamydia at day 8 of pseudopregnancy presented with higher levels of multiple proinflammatory signaling mediators compared to animals challenged at day 10. The strongest differences were observed with IP-10 and IL-5, although significant increases in IL1β CXCL1, IFNγ, IL-6, and IL27p28 were also detected. As these cytokines and chemokines were previously shown to correlate with protection against *C. muridarum* [43–47], this suggested that differences in the proinflammatory cytokine and chemokine response may explain the varying protection observed between day 8 and day 10 of pseudopregnancy.

**Figure 5.**
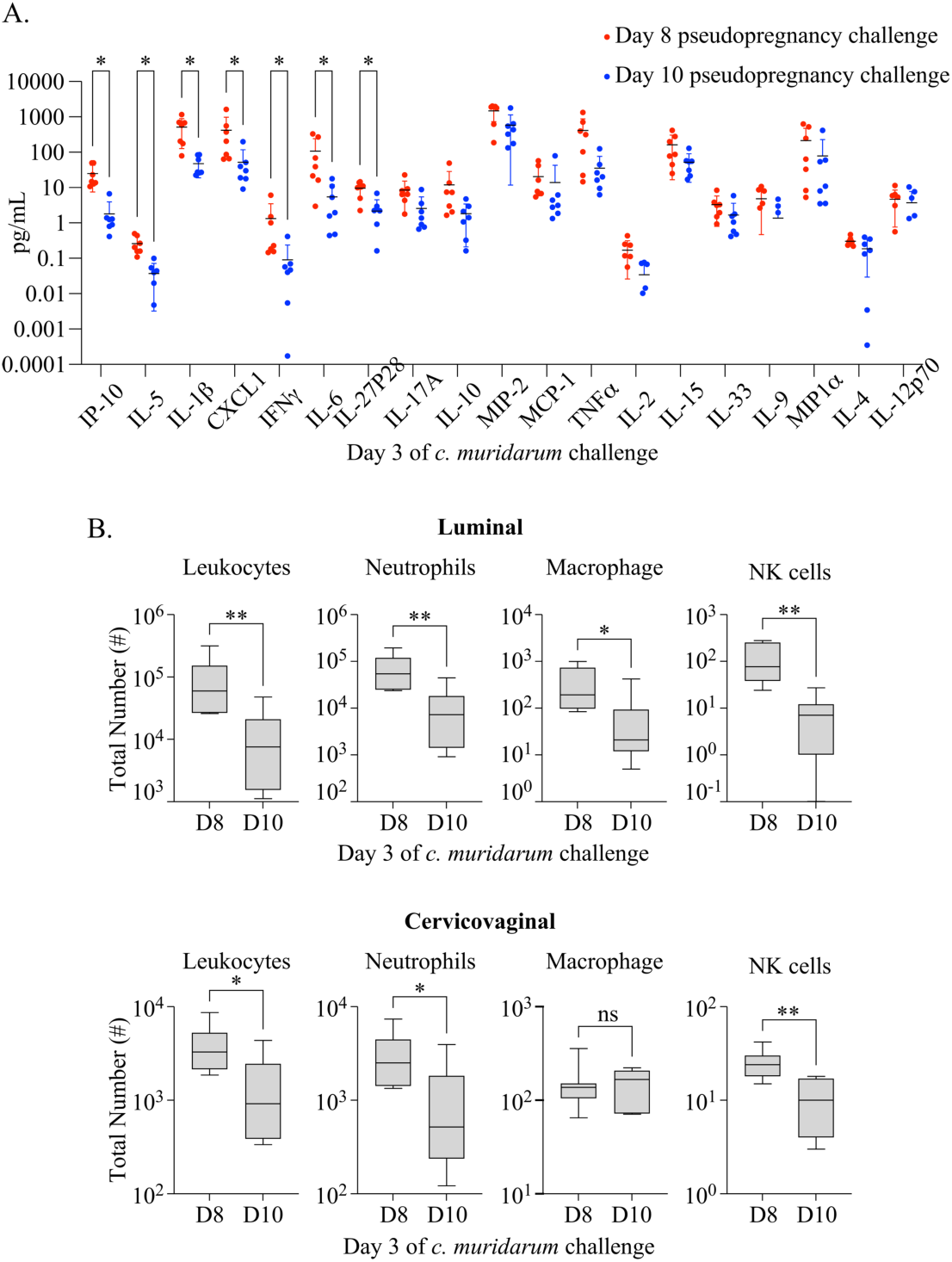
**(A)**. A dot plot graph with the mean and standard deviation (SD) depicting the concentration of indicated proinflammatory cytokine and chemokines measured from CVL supernatant at day 3 of *C. muridarum* infection following challenge at D8 (red) or D10 (blue) of pseudopregnancy. Models used to compare a difference of means were fit using Multiple Mann-Whitney tests and ordered by rank: p-values with q-values ≤ 0.05 are indicated by an asterisk. **(B).** Box and whiskers graphs comparing the total number of indicated IV-negative innate cell populations from CVL (top panels) and vaginal tissues (bottom panels) collected on day 3 of *C. muridarum* infection following challenge at day 8 (D8) or day 10 (D10) of pseudopregnancy. Models used to compare a difference of means were fit using unpaired t-tests. **(A-B).** *p≤0.05, **p<0.01, ***p<0.001, ****p<0.0001

Next, we evaluated the local cellular immune responses at these time points by measuring LFRT and luminal neutrophil, macrophage, and NK cell populations **(Figure 5B)**. These data showed that, in addition to proinflammatory cytokines/chemokines, the numbers of FRT leukocytes were increased in mice challenged on day 8 of pseudopregnancy as compared with day 10. Specifically, the greatest yields and mean differences in overall leukocytes, neutrophils, macrophage, and NK cells were measured from the luminal surface. From the LFRT tissues, these immune populations were also increased following challenge at day 8 as compared with day 10, with the exception of macrophage populations, which were not significantly changed. Taken together, this profiling approach showed there is a more robust early immune response at the infection site in mice challenged on day 8 of pseudopregnancy, suggesting that the immune events that accompany endometrial remodeling are also able to provide a more rapid effector response enabling more effective control of *C. muridarum* infections.

Because LFRT and luminal NK cells were uniquely increased at day 8 of pseudopregnancy, when mice were least likely to exhibit productive *C. muridarum* infections following challenge, we hypothesized that NK cells played a critical role in preventing a productive infection. To test this, we performed antibody-mediated depletion of NK cells over the time points of endometrial remodeling through 2 series of intraperitoneal (IP) injections with αNK1.1 at day 5 and day 7 of pseudopregnancy **(Figure 6A)**. As expected, αNK1.1 administration significantly reduced LFRT and luminal NK cells on day 6 and day 8 of pseudopregnancy **(Figure 6B)**. Next, we vaginally challenged NK cell-depleted mice at day 8 with *C. muridarum* and measured bacterial burden over the infection course **(Figure 6C)**. In contrast to the protection observed at day 8 when NK cells are present **(Figure 4B)**, depletion of NK cells resulted in a productive infection from day 8 challenge, with similar infection kinetics compared to mice administered MPA. Taken together, these data show that the immune events of endometrial remodeling and menstruation lead to NK cell recruitment into the LFRT and cervicovaginal lumen, which plays an essential role in early defense against *C. muridarum*.

**Figure 6.**
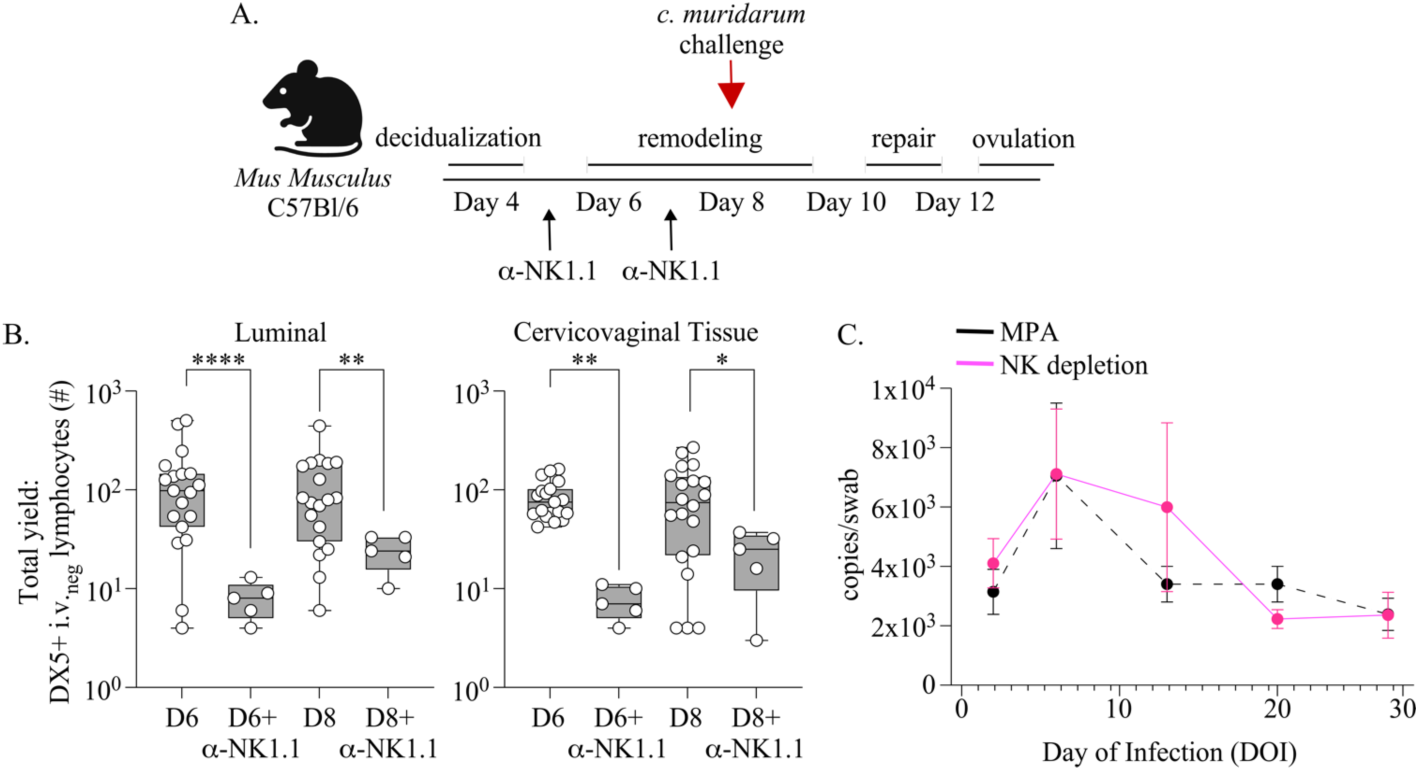
**(A)**. Schematic depicting the approach for depleting NK cells during time points of endometrial remodeling and prior to vaginal *C. muridarum* challenge at day 8 of pseudopregnancy. An IP injection of αNK1.1 antibody is administered on day 5 and day 7 of pseudopregnancy. Schematic created using BioRender.com. **(B).** NK cells are measured at the indicated time points over pseudopregnancy from the cervicovaginal lumen (left panel) or underlying tissues (right panel) following NK cell depletion and compared with mice not treated with αNK1.1 antibody over pseudopregnancy (originally shown in Figure 2). **(C).** The bacterial burden of *C. muridarum* is measured from vaginal swabs collected over the course of infection from mice that are administered αNK1.1 antibody (n=11, pink lines) over endometrial remodeling and compared with historical measurements from mice treated with MPA prior to challenge (n=10, dotted line originally shown in Figure 4B, C). **(B, C).** Models used to compare a difference of means were fit using multiple comparisons: p-values with q-values ≤ 0.05 are shown *p≤0.05, **p<0.01, ***p<0.001, ****p<0.0001.

## Discussion

A growing body of work demonstrates that identifying immune correlates of protection is important for developing biomedical interventions that can prevent infections or limit disease burden caused by pathogens, including those that cause STIs [50–56]. Although uncovering the complex relationships that occur between the immune response and an invading pathogen can provide critical information for understanding how to prevent diseases, our insights into the potential roles of the menstrual cycle in determining such correlates are greatly limited. Despite the fact that the immune system is fundamental for menstruation, elucidating how this process can also impact mucosal immune defense against genitourinary pathogens, including *C. trachomatis,* has been obstructed by the lack of accessible animal models. The experimental and genetic tools available for laboratory mice would allow mechanistic investigations into regional immune dynamics occurring throughout the FRT under menstrual cycle regulation and determination of how changes to immune barrier defenses can alter infection outcomes. Here, we employed the murine pseudopregnancy approach for inducing menstruation in the context of a vaginal chlamydia challenge to explore how the process of endometrial remodeling and repair drives spatiotemporal immune changes in the FRT mucosae and directly test how these alterations shape infections by *C. muridarum*. Using this approach, we discovered that over the course of decidualization, endometrial remodeling, uterine repair, and ovulation, the cervicovaginal tissue undergoes substantial immune alterations that closely mimic the immune changes occurring simultaneously in uterine tissues, and these alterations correlate with protection and infection risk from *C. muridarum* infection. These changes are characterized by an innate immune cell influx of neutrophils, macrophage, and NK cells paired with an increase in local proinflammatory cytokines, particularly IFNγ, during the time frame of endometrial remodeling. The only notable exception we detected from our immune profiling of the lower FRT tissues was the discovery of a sustained population of cervicovaginal NK cells during conditions of progesterone withdrawal when the uterine NK cell populations were contracting. We further confirmed that NK cell fluctuations were also occurring in pig-tailed macaques, a species that naturally undergoes endocrine-controlled menstruation similarly to humans.

From our vaginal *C. muridarum* infection approach over pseudopregnancy, we discovered that challenges administered at decidualization and during endometrial remodeling were correlated with significantly lower bacterial burdens, while challenges administered at menses onset and uterine repair were associated with robust infections. Although we were unable to profile the FRT tissue-localized immune cell populations during menses due to an inability to perform accurate IV labeling of the uterine horns when endometrial vasculature is disrupted, we could still identify a reduction in proinflammatory cytokines and overall leukocyte populations in the cervicovaginal tissues within the broader time frame comprising uterine repair. Interestingly, previous investigations have identified that endometrial repair begins at the very start of decidual shedding and is typified by an immune shift towards more immunosuppressive properties, which are needed to prevent tissue scarring and allow regeneration [43, 44]. In this study, we found that challenges occurring during this time point resulted in robust infections and increased bacterial burdens. Thus, the trajectory of infection initiating during conditions of uterine repair and progressing over the course of endometrial regeneration and ovulation suggests that these specific properties are more hospitable for chlamydial infections in the cervicovaginal tissues. However, future studies will be needed to comprehensively examine this time span in the murine menstruation model in order to better identify and understand those potential risk factors.

Contrary to conditions of uterine repair, *C. muridarum* challenges occurring over decidualization and endometrial remodeling resulted in enhanced protection against infection. As previously described, some of the immune changes correlating to this observation were an increase in local proinflammatory cytokines, especially IFNγ, which is known to be predominantly produced by NK cells. IFNγ signaling has been shown to provide an effective defense against *C. muridarum* in part through inhibiting bacterial replication within infected cells [57–60]. Yet, previous reports have identified that *C. trachomatis*, which naturally infects the FRT in humans via sexual contact, can evade at least some IFNγ-mediated effector functions, which might weaken the relevance of these findings [61, 62]. Notably, we identified that one of the most protective time points over pseudopregnancy was during progesterone withdrawal when IFNγ levels were decreasing, suggesting that IFNγ elevations at the time of challenge were not necessarily essential for early protection. However, both *C. trachomatis* and *C. muridarum* have been shown to be susceptible to NK cell-mediated killing of host cells. Furthermore, NK cells can enhance the activation of Th1 cells, which can provide effective primary and secondary responses against chlamydial infections in the FRT [3, 31, 63–65]. Therefore, to directly test the ability of NK cells to protect against chlamydial infection under conditions of endometrial remodeling, we depleted these cells in mice during this time frame to determine their direct contribution to early protection following *C. muridarum* challenge. These data showed that NK cell-specific depletion resulted in productive infections during endometrial remodeling, similar to the levels of bacterial replication detected in mice administered the hormonal contraceptive MPA, which is most commonly used in murine chlamydia infection models. Thus, these findings demonstrate that NK cell localization at the cervicovaginal mucosa was essential for providing rapid immune protection against primary chlamydial infections.

To better explore how NK cell positioning within the cervicovaginal environment was correlated with protection, we distinguished luminal cells collected by lavage from those embedded into the tissues. Compared with mice on MPA, this approach identified increased NK cell enrichment at the lower FRT luminal barrier during endometrial remodeling. This finding suggests that NK cell proximity to the site of pathogen exposure was important for providing rapid defense that resulted in early protection against chlamydia. Thus, future studies focused on understanding how the menstrual cycle determines NK cell recruitment into the FRT and cervicovaginal lumen might provide greater insights into key chemotactic signals that regulate NK cell localization and could potentially be targeted to enhance immune protection.

The baseline “activation” states of both the innate and adaptive arms of the immune system have been previously shown to predict vaccine efficacies and the likelihood of disease development [56, 66–68]. For example, recent work has identified that basal states of innate immune cells, including NK cell populations, can indicate greater protection against disease development following influenza infection [56]. While many host-intrinsic properties such as age, sex, health, and genetics can influence immune homeostasis, the specific contributions of the menstrual cycle to these baseline immune states in the context of disease risk and vaccine efficacies are far less characterized. Given the role of NK cells in activating adaptive immune responses, including T cells, it is tempting to speculate that the oscillating proinflammatory signals occurring over the menstrual cycle, systemically and within the FRT, might influence both protection from infection and the establishment of immune memory. Thus, this possibility should be further explored in the context of the menstrual cycle

To conclude, this study demonstrates that the process of menstruation regulates regional immune states throughout the upper and lower FRT, which can influence mucosal barrier defenses against chlamydial infections. We further demonstrate that the murine pseudopregnancy approach for inducing menstruation is a valuable tool for investigating how this process drives FRT immune dynamics, and we posit this approach will be beneficial in the development of novel biomedical strategies that can strengthen immunity against genitourinary pathogens.

## Material and Methods

### Mice

C57BL/6J (wild-type) mice and Swiss Webster outbred mice (Taconic Biosciences) were housed under specific Animal Biosafety Level 2 conditions at Emory University. All experiments were performed in accordance with Emory University IACUC guidelines. <u>Pseudopregnancy</u>: 6-12 week old female C57BL/6J were mated with Swiss Webster vasectomized males to induce pseudopregnancy. At 4 days post-mating, female mice received an intrauterine injection of sesame seed oil (Sigma-Aldrich) using mNSET™ devices (Paratechs) as previously described [23]. Cycle phase kinetics were determined using vaginal cytology via Hemotoxin and Eosin staining (H&E), in addition to visible detection of menstruation. For controls, a subset of mice (not undergoing pseudopregnancy) received a subcutaneous injection with 3mg of Medroxyprogesterone acetate (MPA, Prasco) 2 weeks prior to necropsies. <u>NK cell depletion</u>: To deplete NK cells in vivo, mice were intraperitoneally (IP) injected with 200ug α-NK1.1 (clone PK136, BioXcell).

### C. muridarum challenge

*Chlamydia muridarum (C. muridarum)* Mouse Pneumonitis Nigg II strain (ATCC) was cultured in HeLa cells and purified by density centrifugation as previously described [42]. Aliquots were stored in sodium phosphate glutamate buffer (SPG) at -80°C. The inclusion forming units (IFU) from purified elementary bodies were determined by infection of HeLa 229 cells and enumeration of inclusions by microscopy. For vaginal infection, 10^5^ IFU of *C. muridarum* in SPG buffer was deposited into the vaginal vault as previously described [42]. To measure bacterial burden, DNA was purified (Qiagen) from vaginal swabs (Puritan^®^) and quantified by PCR.

### PCR

The *C. muridarum* bacterial burden was measured using Droplet Digital^TM^ PCR (ddPCR^TM^) technology (Bio-Rad) according to manufacturer recommendations [70–72] and was first validated for bacterial burden using *C. muridarum* standards (**Supplemental Figure 2**). In brief, a mixture containing 2x QX200TM ddPCR^TM^ EVAgreen^®^ supermix, mixed 16SR (chlamydia muridarum) forward (AGTCTGCAACTCGACTAC) and reverse (GGCTACCTTGTTACGACT) primers (4µM), ultrapure water, and the DNA sample was used to amplify a fragment of the gene of interest. 20µL of this mixture was added to 70µL of droplet generation oil, and after the droplet generation step, the suspension was used to perform ddPCR in a 96-well PCR plate. The fluorescent signal was read by a QX200^TM^ Droplet Reader (Bio-Rad) and analyzed with QuantaSoft software. The gating for positive droplets was set according to the positive and negative controls read with each plate.

### Murine tissue processing

<u>Intravascular staining:</u> Intravascular staining in mice was performed prior to euthanasia and tissue harvest as previously described[16]. In brief, to discriminate immune cells resident in various tissues from those in circulation, 1.5µg fluorophore-conjugated anti-CD45 Ab in 200µl 1xPBS was IV-injected into the tail vein of mice; 15 minutes post-injection, mice were euthanized with Avertin (2,2,2-tribromoethanol; Sigma-Aldrich) and exsanguinated prior to CVL and tissue collection. <u>CVL collection</u>: To collect and compare cervicovaginal luminal cells in mice, 50ul of sterile PBS was deposited and retracted into the vaginal vault at equal repetitions lasting about 30 seconds. <u>Tissue Processing</u>: FRT tissues were digested using collagenase type II (62.5 U/ml) and DNase I (0.083 U/ml) (STEMCELL Technologies). Cell suspensions were separated by Percoll (GE healthcare life sciences) discontinuous density centrifugation. Enriched leukocytes were washed and resuspended in cell media for phenotyping. For measurement of sex hormones from mice, blood was collected at necropsy by cardiac puncture into 1.3mL EDTA blood tubes (Fisher Scientific) and then centrifuged for plasma collection.

### NHP

For this study, blood and CVL were collected from 6 healthy female pig-tailed macaques of reproductive age over a period of 9 weeks. All NHP procedures were first approved by the CDC Institutional Animal Care and Use Committee. Macaques were housed at the CDC under the full care of CDC veterinarians in accordance with the standards incorporated in the *Guide for the Care and Use of Laboratory Animals* (National Research Council of the National Academies, 2010). All procedures were performed under anesthesia using ketamine, and all efforts were made to minimize suffering, improve housing conditions, and provide enrichment opportunities. 5mL of blood was collected in 8 mL sodium citrate-containing CPT™ tubes (BD Biosciences) and separated into plasma and PBMC by centrifugation. CVL specimens (10 mL collections) were processed as previously described [68, 69].

### Sex-hormone measurement and estimating menstrual cycle phase

Progesterone [P4] and Estradiol [E2] levels in plasma were quantified by immunoassay in one single batch per species. Assay services were provided by the Biomarkers Core Laboratory at the Yerkes National Primate Research Center. The menstrual cycle phase of pig-tailed macaques was estimated by P4 and E2 kinetics relative to a 32-day menstrual cycle (average length of pigtail macaque menstrual cycle) and by observed menstruation.

### Soluble Cytokine/Chemokine Measurement

CVL supernatant and blood plasma were measured and analyzed for cytokine/chemokines through the Emory Multiplexed Immunoassay Core using the Meso Scale Discovery (MSD) platform using a murine and NHP multiplex assay kit in one batch run per species.

### Flow cytometry

Single-cell suspensions were first stained for viability using Zombie NIR^™^ Fixable Viability Kits (Biolegend^®^), followed by cell surface staining and measurements using a BD LSRFortessa™ or LSRII high-parameter cell analyzer, and flow data was acquired using FACS DIVA software (BD Biosciences). Data was analyzed using FlowJo software (TreeStar, Inc.). The following fluorochrome-conjugated antibodies were used:

### Murine antibodies

**Murine antibodies:**
| Marker | Clone | Channel | Company |
| --- | --- | --- | --- |
| IV-CD45 | 30-F11 | BD Horizon™ PE-CF594 | BD Biosciences |
| CD45 | 30-F11 | BD Pharmingen™ FITC | BD Biosciences |
| CD49b | DX5 | Phycoerythrin (PE) | BioLegend® |
| CD68 | FA-11 | PerCP/Cyanine5.5 | BioLegend® |
| F4/80 | BM8 | Brilliant Violet 650™ | BioLegend® |
| CD11b | M1/70 | BD Horizon™ BUV395 | BD Biosciences |
| MHCII | M5/114.15.2 | PE/Cyanine7 | BioLegend® |
| Ly-6G | 1A8 | Brilliant Violet 510™ | BioLegend® |
| CD115 | T38-320 | BD OptiBuild™ BUV496 | BD Biosciences |

### NHP antibodies

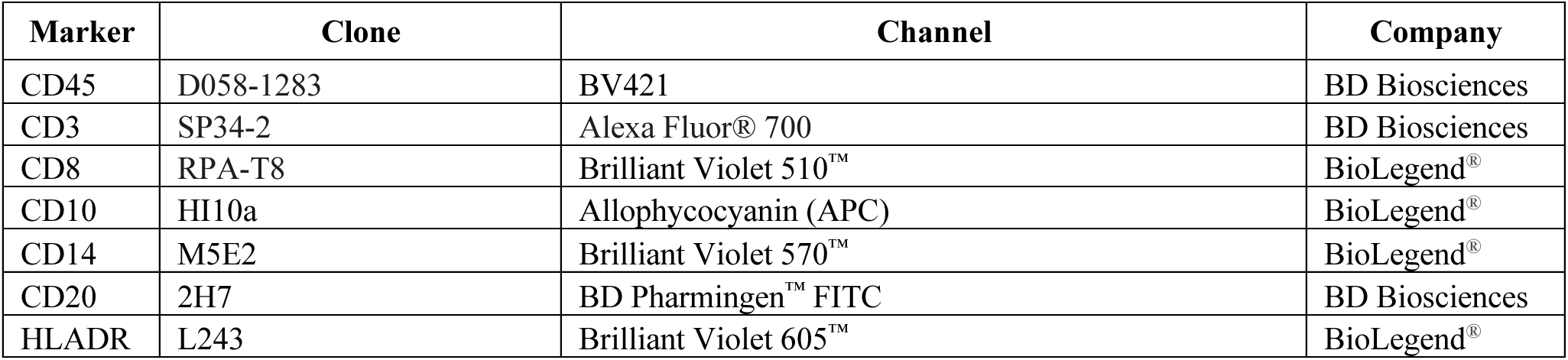

### Funding

This study was supported by NIH grant R21AI180610 (A.S-K).

(sex hormone levels) Assay services were provided by the Biomarkers Core Laboratory at the Emory National Primate Research Center. This facility is supported by the Emory National Primate Research Center Base Grant P51 OD011132.

(Cytokines) This study was supported in part by the Emory Multiplexed Immunoassay Core (EMIC), which is subsidized by the Emory University School of Medicine and is one of the Emory Integrated Core Facilities. Additional support was provided by the National Center for Georgia Clinical & Translational Science Alliance of the National Institutes of Health under Award Number UL1TR002378.

W.M.S. is a recipient of a Senior Research Career Scientist from the Medical Research Service of the Department of Veterans Affairs.

The content is solely the responsibility of the authors and does not necessarily reflect the official views of the National Institutes of Health. (RRID:SCR_023528), the Centers for Disease Control and Prevention, or the Department of Veterans Affairs.

## Acknowledgments

We thank Dr. Marion Rudolph at Bayer HealthCare for helpful insights. From the CDC Division of HIV Prevention (DHP), we thank Dr. J. Gerardo-Garcia Lerma, Dr. Jim Smith, Sunita Sharma, Susan Rhone, James Mitchell, and Frank Deyonks for their assistance in the NHP studies. From Emory University Medical School, we thank Dr. Jacob E. Kohlmeier for assistance with mouse studies.

## Supplemental Data

**Supplemental Figure 1:**
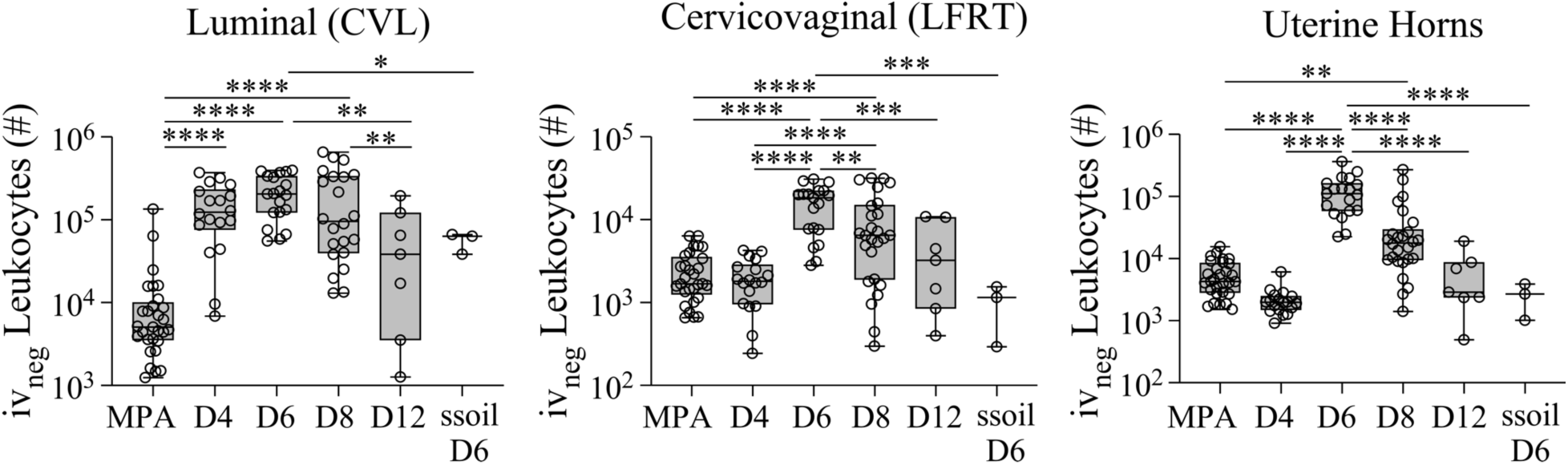
The total leukocyte yield from indicated FRT tissue sites is plotted as bar and whiskers graphs over pseudopregnancy and compared with mice administered MPA or mice administered sesame seed oil in the absence of pseudopregnancy as a control. Models used to compare a difference of means were fit using multiple comparisons: p-values with q-values≤0.05 are shown *p≤0.05, **p<0.01, ***p<0.001, ****p<0.0001.

**Supplemental Figure 2:**
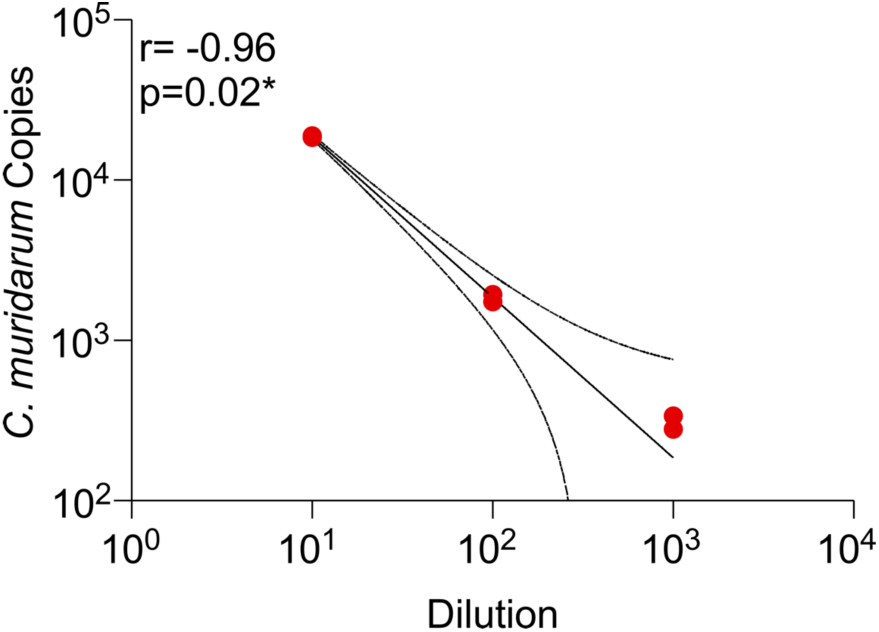
An XY graph with prediction bands plotting the ddPCR quantification using dilutions taken from DNA extracted from 1x10^5^ IFU of *C. muridarum*. Distributions were tested by Spearman’s correlations.

**Supplemental Table 1:** the mean levels of indicated cytokines and chemokines (pg/mL) with the SEM over pseudopregnancy and in mice administered MPA.

|  | **MPA (pg/mL)** | **Day 4 (pg/mL)** | **Day 6 (pg/mL)** | **Day 8 (pg/mL)** | **Day 10 (pg/mL)** | **Day 12 (pg/mL)** |
| --- | --- | --- | --- | --- | --- | --- |
| *IL15* | 18.825 (3.489) | 96.336 (36.655) | 218.936 (22.966) | 90.755 (24.473) | 181.386 (47.04) | 32.26 (14.009) |
| *IL-17A* | 1.242 (0.185) | 9.915 (2.939) | 80.811 (12.351) | 9.481 (2.765) | 10.151 (3.025) | 2.633 (0.822) |
| *IL-27P28* | 1.658 (0.421) | 4.571 (1.316) | 26.045 (4.042) | 12.289 (5.05) | 16.329 (5.771) | 2.016 (0.623) |
| *IL-33* | 13.544 (4.126) | 7.508 (3.059) | 5.522 (0.594) | 11.091 (3.077) | 9.911 (3.03) | 1.881 (0.717) |
| *IL-9* | 0.873 (0.572) | 1.856 (.0878) | 4.096 (1.016) | 1.587 (0.863) | 3.234 (2.156) | 1 (0.521) |
| *IP-10* | 8.959 (1.146) | 15.375 (4.145) | 845.045 (216.52) | 103.941 (33.999) | 119.646 (49.228) | 5.081 (1.767) |
| *MCP-1* | 2.056 (0.4) | 8.631 (4.181) | 17.370 (2.771) | 19.829 (5.712) | 68.625 (42.208) | 53.521 (39.849) |
| *MIP-1α* | 12.803 (1.441) | 169.677 (65.429) | 329.283 (54.306) | 147.607 (67.654) | 253.184 (103.611) | 24.035 (13.966) |
| *MIP-2* | 132.679 (21.303) | 1188.208 (244.673) | 1936 (24.36) | 997.099 (250.718) | 1461.614 (250.464) | 530.557 (202.535) |
| *IFNγ* | 0.049 (0.012) | 0.2 (0.134) | 104.177 (20.15) | 1.691 (0.599) | 16.806 (13.003) | 0.201 (0.114) |
| *IL-10* | 0.733 (0.223) | 16.69 (8.08) | 64.891 (14.663) | 22.646 (9.159) | 30.386 (14.889) | 0.882 (0.258) |
| *IL12p70* | 1.339 (0.494) | 3.596 (1.177) | 30.338 (4.65) | 7.966 (3.07) | 13.117 (4.122) | 1.753 (0.788) |
| *IL-1β* | 393.715 (55.892) | 285.274 (75.729) | 1341.195 (174.339) | 1163.974 (695.134) | 1665.124 (1178.348) | 157.367 (63.492) |
| *IL-2* | 0.038 (0.012) | 0.071 (0.017) | 0.766 (0.13) | 0.599 (0.219) | 0.57 (0.253) | 0.063 (0.03) |
| *IL-4* | 0.169 (0.033) | 0.089 (0.025) | 0.495 (0.092) | 0.39 (0.131) | 0.249 (0.048) | 0.216 (0.076) |
| *IL-5* | 0.389 (0.056) | 0.615 (0.219) | 1.867 (0.492) | 1.356 (0.527) | 1.071 (0.373) | 0.14 (0.035) |
| *IL-6* | 2.16 (0.388) | 42.375 (14.848) | 7741.421 (1358.467) | 714.485 (250.752) | 2950.566 (1589.784) | 9.655 (5.112) |
| *CXCL1* | 62.722 (14.326) | 137.711 (86.099) | 846.746 (138.84) | 519.134 (188.805) | 882.652 (267.625) | 37.582 (12.072) |
| *TNFα* | 8.714 (0.987) | 295.473 (82.585) | 2882.011 (357.084) | 367.174 (148.467) | 371.82 (127.41) | 49.419 (20.185) |

